# Sister chromatid cohesion halts DNA loop expansion

**DOI:** 10.1101/2023.07.31.551217

**Authors:** Nathalie Bastié, Christophe Chapard, Sanae Nejmi, Henri Mboumba, Agnès Thierry, Frederic Beckouët, Romain Koszul

**Author notes:** ^&^ these authors contributed equally to this work (Nathalie Bastié, Christophe Chapard).

## Abstract

Eukaryotic genomes are folded into DNA loops mediated by structural maintenance of chromosomes (SMC) complexes such as cohesin, condensin and Smc5/6. This organization regulates different DNA-related processes along the cell cycle such as transcription, recombination, segregation and DNA repair. During G2/M stages, SMC-mediated DNA loops coexist with cohesin complexes involved in sister chromatid cohesion (SCC). However, the articulation between the establishment of SCC and the formation of SMC-mediated DNA loops along the chromatin remains unknown. Here we show that SCC is indeed a barrier to cohesin-mediated DNA loop expansion in G2.

## Instroduction

Eukaryotic genomes are folded into chromatin loops and self-interacting domains by structural maintenance of chromosome (SMC) complexes (cohesin, condensin and Smc5/6). This organisation regulates or influences various DNA-related processes throughout the cell cycle, such as transcription, recombination, DNA segregation and repair ^1–3^. During G2/M phases, SMC complexes mediating DNA loops coexist with the cohesin complex, which is long known for its involvement in maintaining sister chromatid cohesion (SCC). Whether and how SCC and DNA looping influence each other remains unclear (Davidson and Peters 2021).

Cohesin is composed of two Smc proteins (Smc1 and Smc3) and one α-kleisin subunit, Scc1/Rad21 ^4^. Both Smc proteins are rod-shaped and form long antiparallel coiled coils. They dimerize via their hinge domain at one extremity and are linked at the other end by ATPase head domains that interact with the kleisin subunit ^5^. Several studies in budding yeast suggest that cohesin mediates SCC by topologically embracing the two sister chromatids within compartments formed within the cohesin ring complex ^6–8^. In this way, the two sisters are held together from DNA replication to their segregation during mitosis, ensuring the correct transmission of the genetic information.

Loading of cohesin onto chromosomes depends on a loading factor called Scc2/NIPBL ^9^, which also has the ability to stimulate SMC ATPase activity in the presence of DNA. In contrast, the “releasing activity”, which removes cohesin from DNA, allows dynamic association with chromosomes, particularly in the G1 phase of the cell cycle ^10, 11^. The later relies on two cohesin-associated regulatory subunits called Wpl1/WAPL and Pds5 ^11^. During S phase, the DNA replication machinery promotes acetylation of two conserved Smc3 lysines (K112 and K113) by the Eco1/ESCO1 acetyltransferase to inhibit the “releasing activity” and, consequently, establish stable sister chromatid cohesion. Smc3 acetylation is then maintained during G2/M and is only removed during anaphase, when the kleisin subunit is cleaved by Separase ^12–15^.

*In vivo* and *in vitro* experiments have recently revealed that human cohesin can establish and enlarge chromatin loops along DNA molecules ^16–19^. These loops are important for partitioning human interphase chromosomes into domains and regulating processes such as transcription, DNA recombination or repair ^3, 20, 21^. It has been proposed that cohesin captures small DNA loops and then moves DNA using an internal molecular motor to enlarge the loops processively ^22, 23^. This process, known as the loop extrusion model, also organizes into loops the chromosomes of budding yeast *Saccharomyces cerevisiae* during the G2/M stages of the cell cycle ^24, 25^. In yeast, these loops are controlled by a number of conserved mechanisms. First, Wpl1-mediated release limits DNA loop size ^24, 25^. Second, Scc2-stimulated ATPase activity is required for DNA loops expansio ^16, 26, 27^. Third, by competing with Scc2 for binding to cohesin, Pds5 inhibits DNA loops expansion. Finally, Eco1-mediated Smc3 acetylation negatively regulates loop expansion ^26–29^. However, how Smc3 acetylation regulates loop expansion remains unknown. A possibility is that acetylation of cohesin-extruding DNA loops along each sister maintains these loops at punctate positions along chromosomes. Another is that the acetylated cohesins that stably entrap sister chromatids represent physical barriers to DNA extruding cohesins. To understand how different pools of cohesin and their associated dynamics contribute to DNA loop formation, we analyzed how wild-type cohesins involved in cohesion regulate the expansion of DNA loops along sister chromatids by a cohesin mutant (*scc1-V137K*) which is defective for Pds5 binding ^30^. We show that this mutant mimics the previously described effects of Psd5 depletion and further demonstrate that acetylated wild-type cohesin is a non-permissive barrier to *scc1-V137K* mediated DNA loop expansion. Our results reveal for the first time that different cohesin complexes cooperate to regulate the positioning of DNA loops along chromosomes.

## Results

To analyze whether cohesin complexes influence each other in their ability to expand DNA loops, we explored how the spreading of intra-chromosomal contacts to unusual long distances by a mutant form of cohesin, that cannot form cohesion, can be impacted by a pool of wild-type cohesin maintaining SCC.

### scc1-V137K spread intrachomosomal contact over great distances

In the absence of Pds5, a cohesin subunit essential for SCC, cohesin is able to expand very long DNA loops (Dauban et al., 2020). It was therefore hypothesized that the Scc1-V137K mutant, defective in both Pds5 binding and SCC, could be an ideal super extruder ^24, 30^. To determine this, we examined the effect of Scc1-V137K on genome organization using a haploid strain carrying both an endogenous auxin-degradable *SCC1* (Scc1-AID) and an ectopic *scc1-V137K* alleles, the expression of which can be controlled by the inducible Gal1-10 promoter. Note that the endogenous *SCC1* expression also ensures cell viability of the otherwise lethal *scc1-V137K* gene. Scc1-V137K protein was induced in G1 synchronized cells, while the endogenous Scc1-AID was concomitantly depleted. The cells were then released and arrested in G2 using nocodazole (**Fig. 1a, 1b, and 1c**), and Scc1-V137K deposition quantified using calibrated ChIP-seq (**Methods**). As expected, Gal induction of *scc1-V137K* resulted in both an enrichment in cohesin at the core CEN, and a depletion of cohesin peaks located on the chromosome arms (**Fig. 1d upper panel**), consistent with the cohesin profile obtained when cells are depleted of Pds5 ^31^. Calibrated ChIP-seq also show that the amount of cohesin between the peaks is higher than that observed for the wild-type condition (**Fig. 1d lower panel**) suggesting that cohesin may slide along the chromatin.

**Figure 1:**
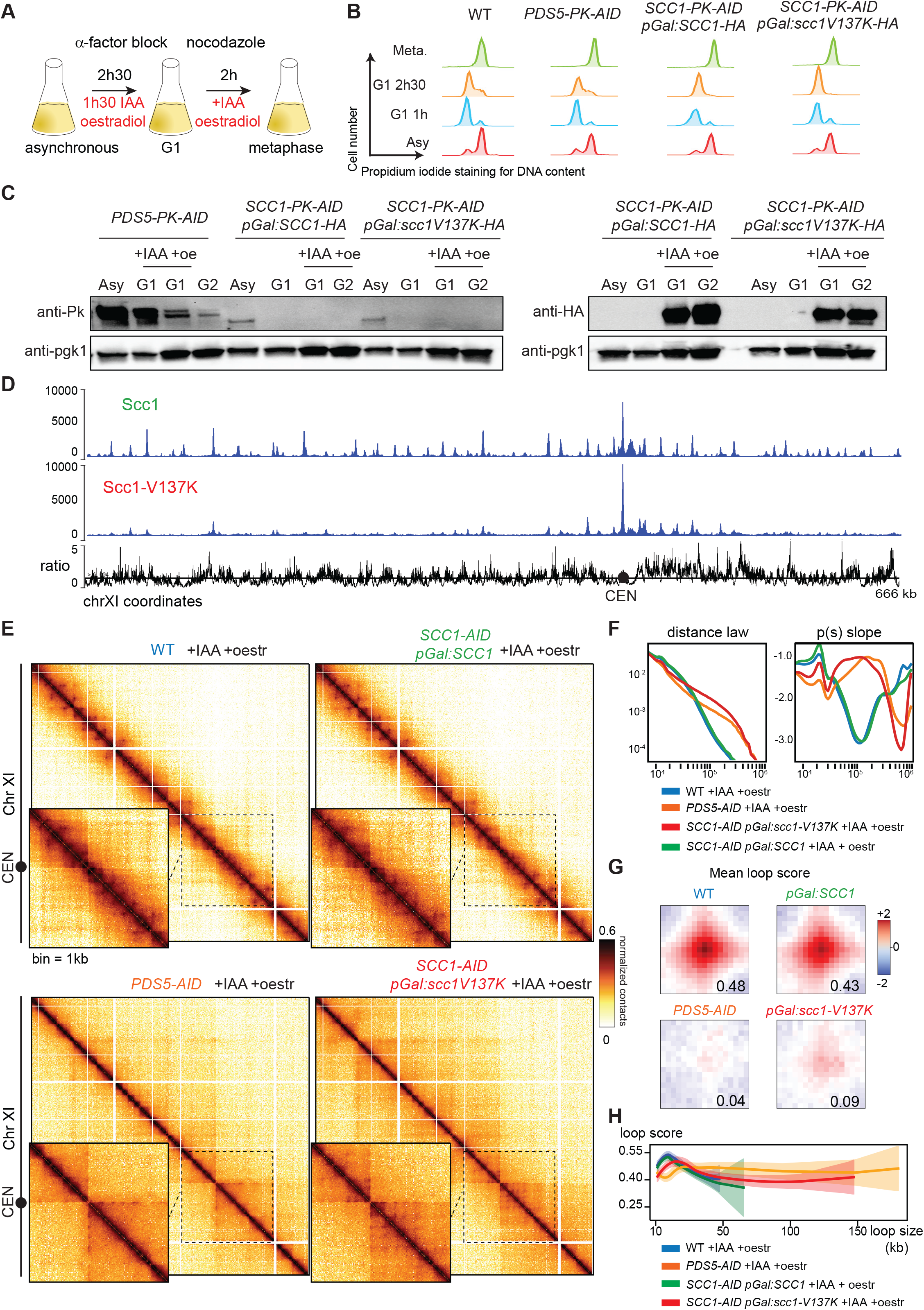
*scc1-V137K* extends loops over long distances. **A)** Experimental protocol used to process synchronized yeast cells (W303-1A, yNB33.1-8a, yNB77.1-2b and yNB78.1-15c) from G1 into metaphase while expressing *scc1-V137K* and SCC1 under Gal1 promoter (+ OE), and in absence of endogenous SCC1 (+IAA). **B)** Cell synchronization monitored by flow cytometry. **C)** Western blot assessing (Left Panel) the loss of Pds5-PK-AID or Scc1-PK-AID, and (Right Panel) the induction of Scc1 or Scc1-V137K under Gal1 promoter (oestradiol induction). Pgk1: loading control. **D)** Calibrated ChIP-seq profiles showing the distribution of Scc1-HA, Scc1-V137K, as well as the ratio between Scc1-HA and Scc1-V137K deposition profiles. Black line: ratio 1. **E)** Hi-C contact maps of chromosome XI (bin: 1kb) from metaphase arrested cells for (top left) WT (W303-1A) and (bottom left) Pds5 depleted cells, and of cells depleted for endogenous Scc1, but expressing (top right) Scc1 or (bottom right) Scc1-V137K under the control of an inducible promoter. A magnification of chromatin contacts within a region of chromosome XI including the centromere (CEN, black dot) are presented. **F)** Left: contact probability curves p(s) representing the average contact frequency as a function of genomic distance (bp). Right: corresponding local derivative curves. **G)** Mean profile heatmap of loops called using Chromosight. **H)** Loop spectrum generated using Chromosight, indicating average loop scores as a function of loop sizes.

To further examine the effect of Scc1-V137K on chromosome organization, we generated Hi-C contact maps for cells expressing either *scc1-V137K* or *SCC1* genes from the Gal1-10 promoter (**Fig. 1a, 1b, and 1c**), wild type cells or Pds5 depleted cells, all processed similarly. The contact maps and probability curves (*p(s)*) showed that Scc1-V137K has comparable (slightly higher) effects than Pds5 depletion. In other words, Scc1-V137K promotes intra-chromosomal interactions over longer distances than those observed in wild-type cells (**Fig. 1e, 1f**). Chromatin loops along chromosomes, reflected by dots on the Hi-C maps, were called using the program Chromosight on subsampled Hi-C maps (**Fig. 1g; Methods)** ^32^. The number of called loops along chromosome arms in the presence of Scc1-V137K (191 loops) is strongly reduced compared to WT cells (523 loops), or to a control strain expressing an ectopic copy of *SCC1* (pGal:SCC1-HA) (507 loops). This number is close to the number of loops called in the absence of Pds5 in a former work (247 loops^26^. In addition, Scc1-V137K differed slightly from Pds5 depletion in the immediate vicinity of the CEN, as grid-like, punctate DNA loops were observed on Hi-C maps. However, comparison of mean loop scores (MLS; **Methods**) confirmed a global reduction in loop signal in cells expressing Scc1-V137K (MLS = 0.09) similar to cells depleted for Pds5 (0.09±0.04; n=3), to compare with SCC1 (0.43) or WT (0.43±0.04; n=4) loops (**Fig. 1g**). In addition, the few remaining DNA loops detected have their basis separated by longer distances (over 100 kb in the presence of Scc1-V137K) compared to control cells (< 50kb), as shown by the loop size distribution spectrum (**Fig. 1h**). Altogether, these experiments show that Scc1-V137K extends intra-chromosomal contacts to a greater extent than wild type, mimicking the effect of Pds5 depletion.

### Cohesin acts as non-permissive barrier to loop extrusion

We then investigated how wild-type cohesins involved in sister chromatids cohesion influence the ability of Scc1-V137K to expand DNA loops. To do this, *scc1-V137K allele* was expressed for 90 minutes in nocodazole-arrested G2 cells that had either been depleted of endogenous Scc1 or allowed to express a wild-type form of Scc1 throughout replication, thereby establishing cohesion (**Fig. 2a,b; Methods**). Western blot and microscopy analysis showed that Scc1-V137K is expressed at similar levels regardless of cell line and did not affect pre-established sister chromatid cohesion (**Fig. 2c, Sup 1a**). Hi-C contact matrices and contact probability curves (*p(s)*) generated from these cells showed that while the expression of Scc1-V137K in G2 cells lacking the wild-type form of cohesin allowed expansion of intra-chromosomal contacts over longer distances to the same extent as when expressed from G1 arrested cells (**Fig. 2d upper panels, 2e**), the expansion of DNA loops by Scc1-V137K was greatly reduced in cells harboring chromosome-bound wild-type cohesin (**Fig. 2d middle panels, 2e and Fig sup. 1b**). Note that Scc1-V137K extrusion activity was not abolished as the compaction level mediated by expression of Scc1-V137K in the presence of WT was slightly higher than in controls cells expressing Scc1-WT (**Fig. 2d lower panels, 2e and Fig sup. 1b**). In addition, chromosome arms formed stronger domains on either side of the centromeres when Scc1-V137K was expressed even in the presence of Scc1-WT, as expected from the absence of Pds5 binding. In addition, and in contrast to WT Scc1 or Scc1-V137K taken separately, expression of Scc1-V137K in the presence of WT cohesin led to the appearance of new punctate DNA loops bringing stable DNA loops together, not only immediately around the CEN but also on the arms. Not only the number of loops were increased in this condition (from 523 to 690) but also the mean loop score (0.50; Fig. **2f**). In agreement, the loop size distribution spectrum shows that detected DNA loops when Scc1-V137K is expressed in the presence of WT cohesin have an intermediate size compared to the smaller WT loops and larger Scc1-V137K loops (**fig. 2g**). Importantly, analysis of the genome wide distribution of Scc1-V137K by calibrated ChIP-seq experiment confirmed that Scc1-V13K loading on DNA was also not abolished by the preloaded wild-type cohesin (**Fig. 2h**). Nevertheless, the presence of wild-type cohesin decreased the amount of Scc1-V137K in between the peaks as compared to Scc1-V137K distribution in the absence of Scc1-WT, reinforcing the idea that wild-type cohesin may inhibit the translocation of Scc1-V137K that mediates long distance DNA loop expansion rather than of other translocating cohesin along the arms. Taken together, our results suggest that DNA loop expansion by the super extruder is impaired or halted by the presence of wild-type cohesin.

**Figure 2:**
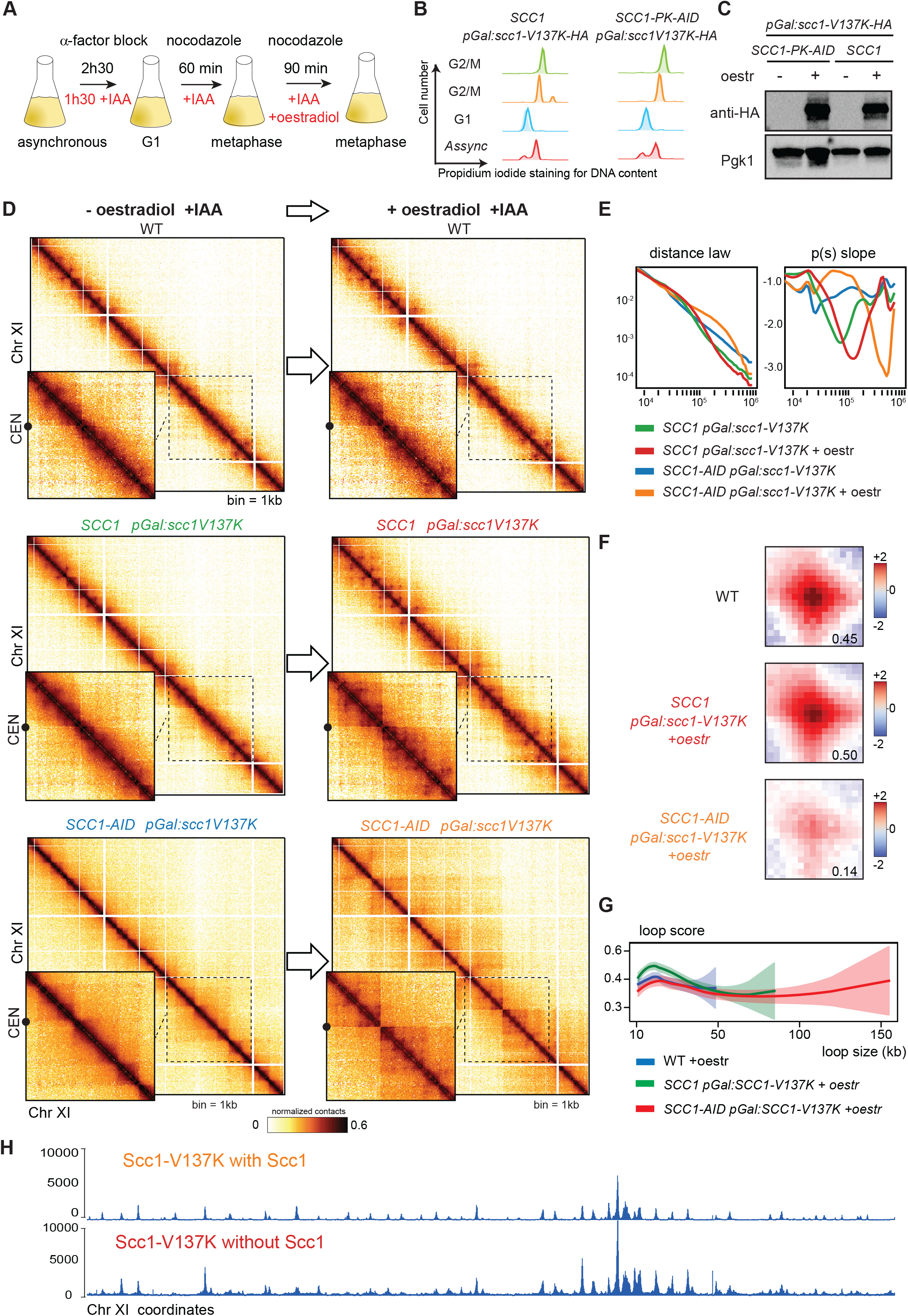
*scc1-V137K*-mediated loop expansion is inhibited by the presence of pre-loaded wild-type cohesin (see also Figure S1) **A)** Experimental protocol used to process yeast cells from G1 to metaphase to express *scc1-V137K* in G2 in absence or in presence of endogenous SCC1. **B)** Cell synchronisation monitored by flow cytometry. **C)** Western blot assessing the induction and expression of *scc1-V137K*. Pgk1: loading control. **D)** Hi-C contact maps of chromosome XI (bin: 1kb) for three nocodazole-arrested strains (WT, *SCC1 pGal:scc1V137K*, and *SCC1-AID pGal:scc1V137K*) synchronized in G2/M in presence of auxin (+IAA), and without (left column) and with (right column) oestradiol addition. Top row: WT control. Middle row: cells expressing Scc1-V137K in presence of endogenous Scc1. Bottom row: cells expressing Scc1-V137K in absence of endogenous Scc1. A magnification of chromatin contacts within a region of chromosome XI including the centromere (CEN, black dot) are presented. **F**) Left: contact probability curves p(s) representing the average contact frequency as a function of genomic distance (bp) of the corresponding strains. Right: corresponding local derivative curves. **F)** Mean profile heatmap of loops called using Chromosight. **G)** Loop spectrum generated using Chromosight, indicating average loop scores as a function of loop sizes. **H)** Calibrated ChIP-seq profiles showing the effect of presence or absence of endogenous SCC1 on Scc1*-*V137K deposition.

### Sister chromatid cohesion is a non-permissive barrier

Since there are at least two populations of WT cohesin bound to mitotic chromosomes, one engaged in sister chromatid cohesion and one involved in DNA looping, we wondered whether sister chromatid cohesion could be the barrier preventing the expansion of DNA loops. If so, preventing the appearance of the “cohesive” cohesin population involved in cohesion should remove or attenuate this inhibition. In absence of Cdc45, G1 synchronized cells can enter G2/M without replication or Smc3 acetylation: in these cells, the unreplicated chromosomes thus lack cohesion, but are nevertheless folded into loops by cohesins. We therefore took advantage of this genetic system to explore the extent to which established wild-type cohesin-mediated loops inhibits Scc1-V137K mediated loop expansion in these cells. We first verified that *scc1-V137K* can form long-range DNA loops in this context. Expression of *scc1-V137K* for 90 minutes in Cdc45-depleted, G2/M-arrested cells also depleted of endogenous Scc1 (**Fig. 3a, 3b and c; Methods**) resulted in an enhanced extrusion activity (**Fig. 3d, 3e**), with fewer (205) but longer DNA loops than in a WT control (**Fig. 3d-f**, ref NSMB). In presence of WT cohesin folding the G2/M, unreplicated chromosomes into loops, the contact maps and associated P(s) of cells expressing *scc1-V137K* still show an increase in long-range DNA contacts compared to regular G2/M replicated cells, although to a lesser extent than in the absence of WT cohesin (**Fig. 3d, 3e**) (**Fig. 2c, 2d, 3c, and 3d**). Therefore, loops mediated by WT cohesins have a moderate inhibitory effect on the expansion of loops mediated by *scc1-V137K*, as compared to G2 cells (**Fig. 3c, d, and Fig sup. 1g)**. Also contrasting with G2 (**Fig. 2**), both the number of DNA loops and associated mean loop score were not increased in Cdc45-arrested cells after induction of *scc1-V137K* in the presence of WT cohesin but were instead attenuated (200 loops after induction as compared to 232 before, MLS=0.27 after induction as compared to 0.40 before; **Fig. 3e**). This revealed that Scc1V-137K can establish longer loops in the presence of cohesin that dynamically associates with chromosomes as compared to the effect observed in the presence of stably bound cohesin after DNA replication. Altogether, our results show that the latter cohesin population, making SCC, represents a non-permissive barrier to DNA loop expansion while while un-acetylated bound WT cohesin is a less efficient barrier than SCC.

**Figure 3:**
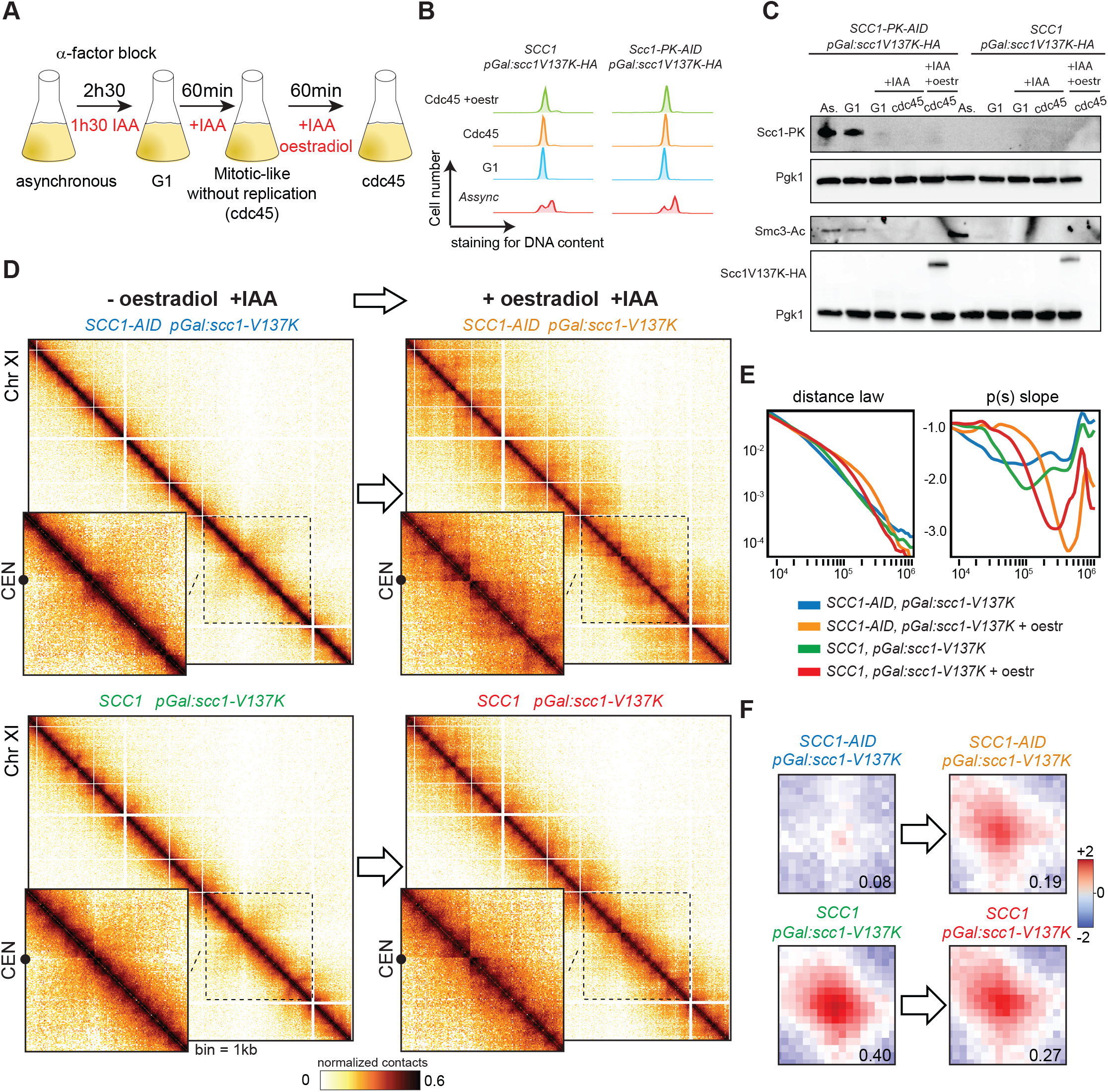
Replication reinforces inhibition of chromatin loop expansion (see also Figure S1) **A)** Experimental protocol used to process yeast cells from G1 into G2/metaphase without replication by depleting Cdc45-AID using auxin (IAA), and then inducing the super extruder *Scc1-V137K* in G2 in absence or in presence of endogenous SCC1. **B)** Cell synchronization monitored by flow cytometry. **C)** Western blot assessing Scc1-PK degradation and *scc1V137K*-HA expression. That Cdc45-AID is depleted was monitored through the absence of Smc3-Ac. Pgk1: loading control. **D)** Hi-C contact maps of chromosome XI (bin: 1kb) for two nocodazole-arrested, unreplicated (cdc45) strains (*SCC1-AID pGal:scc1V137K*, and *SCC1 pGal:scc1V137K*) synchronized in G2/M in presence of auxin (+IAA), and without (left column) and with (right column) oestradiol addition. Top row: unreplicated G2/M cell expressing Scc1-V137K in absence of endogenous Scc1. Bottom row: unreplicated G2/M cells expressing Scc1-V137K in presence of endogenous Scc1. A magnification of chromatin contacts within a region of chromosome XI including the centromere (CEN, black dot) are presented. **E)** Left: contact probability curves p(s) representing the average contact frequency as a function of genomic distance (bp) of the corresponding strains. Right: corresponding local derivative curves. **F)** Mean profile heatmap of loops called using Chromosight.

## Discussion

Loop extrusion is thought to progressively enlarge DNA loops until cohesin encounters a roadblock and/or is released from DNA by Wpl1 ^18, 19^. In mammals, CTCF was originally identified as a major barrier to LE ^17, 33^. However, the bases of DNA loops do not always correspond to CTCF-bound sites, and cohesin can also establish stable loops along chromosomes of organisms that do not possess CTCF ^24^.Thus, other DNA-binding proteins and/or DNA-related processes can prevent loop extrusion. Recently, conserved factors in the replication machinery have been shown to arrest cohesin-mediated DNA expansion in mammals ^34^. In addition, transcriptional activation influences DNA loop expansion in different organisms (Chapard et. al in prep) ^35, 36^. Cohesins actively extruding DNA are also confronted with other cohesins involved in SCC, or other SMCs engaged in other DNA looping process. How these different populations of SMC complexes influence DNA loop extrusion remained poorly understood. For example, they may act as permissive or non-permissive barriers depending on whether they are easily crossed by loop expansion SMCs. *In vitro*, condensin and cohesin can cross each other to form Z-loops ^37^. The present work shows that DNA loop expansion by the super-extruder *scc1V137K* is strongly inhibited *in vivo* by preloaded wild-type cohesins that provide stable sister-chromatid cohesion. Inhibition by the super-extruder also occurs, though to a lower extent, when cells are into a mitosis-like state without replication or Smc3 acetylation, likely reflecting the influence of SMC involved in mitotic loops and compatible with a Z-loop formation (**Figure 4**).

**Figure 4:**
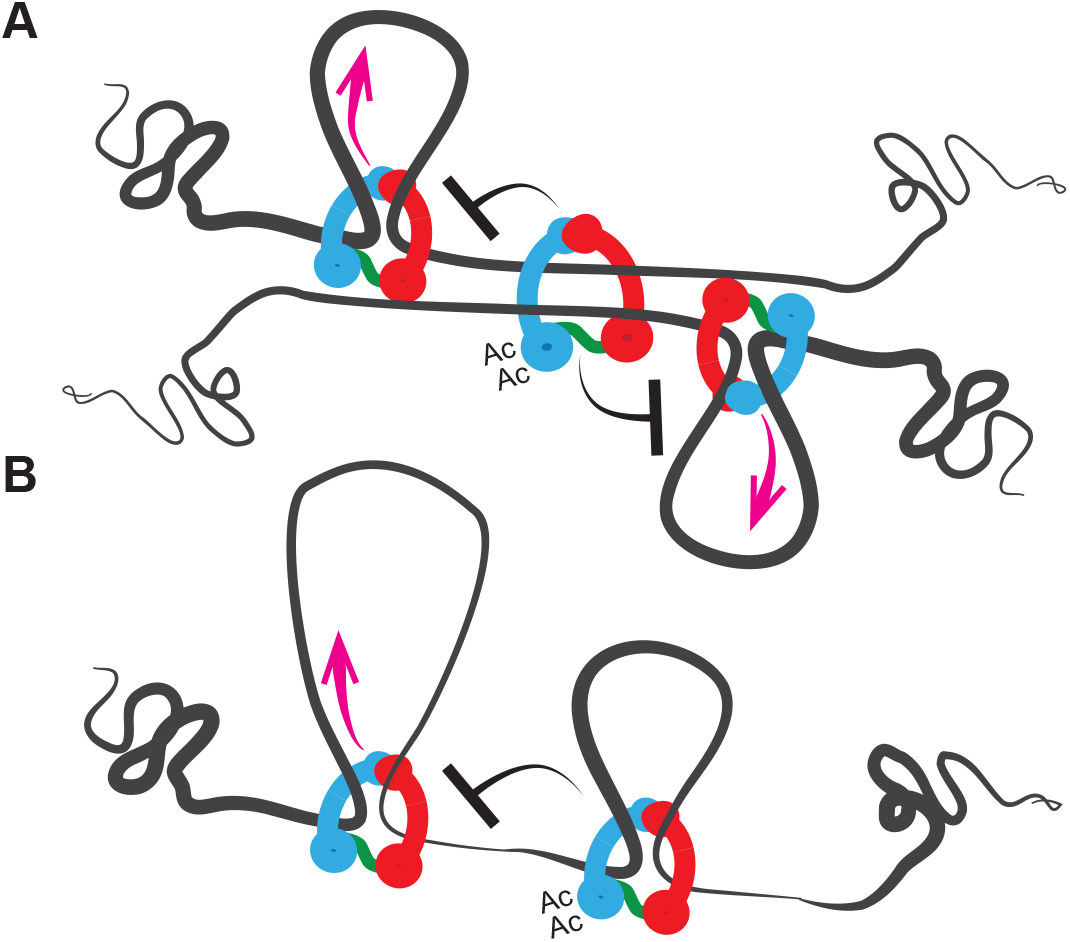
Models for inhibition of chromatin loop extrusion by stable cohesins. **A)** Cohesive cohesin inhibiting loop extrusion **B)** Extrusive cohesin inhibiting loop expansion

As acetylation is involved in sister chromatid cohesion, our data suggest that the pool of cohesin which is committed to SCC could the strongest inhibits loop extrusion in G2 cells while cohesin that does not mediate SCC may only represent a semi-permissive barrier. The acetylation cycle of cohesin is known to regulate DNA loop expansion in yeast as well as in mammals ^26, 27, 38^. A recent study further revealed that Eco1-mediated Smc3 acetylation also inhibits loop expansion on un-replicated DNA in mammalian cells ^27^. It is therefore conceivable that acetylation itself, whether of cohesin promoting SCC or DNA loops, may stop the expansion of DNA loops mediated by non-acetylated cohesin or directly contribute to its process, for instance by stabilizing other proteins or cohesin subunits such as Pds5 ^26^. As experiments in budding yeast have in addition revealed that cohesin can topologically entrap either one or two DNA molecules, other cohesin mediated topological constraints could also represent a barrier for LE on either replicated or non-replicated DNA.

Our result is also supported by the fact that cleavage of Rec8, the meiotic-specific Scc1 analogue, in oocyte increases the length of extruded loop by Rad21 ^39^. If SCC is a barrier to cohesin-mediated loop extrusion, it is likely to be in addition a barrier to other SMC complexes that extrude DNA, i.e. Smc5/6 or condensin. In other words, SCC may also influence condensin-mediated chromosome condensation or Smc5/6-mediated DNA repair. It could also influence the cohesin-mediated reorganization of broken chromosome extremities reported in yeast ^40^.

Two recent studies in mammals’ cells revealed an interplay between cohesin mediated loops and replication machinery ^34, 41^. One revealed that MCM complexes could act as boundaries to halt cohesin mediated loop extrusion and the other one showed that cohesin-mediated loop anchor is an essential determinant of the locations of replication origins. It may be that the interaction between MCM and LE leads to the conversion of DNA loop anchoring cohesin into cohesin engaged in sister chromatid cohesion after passage of the replication fork. Thus, the cohesin involved in sister chromatid cohesion would possibly be located at the basis of preexisting DNA loops.

Overall, our study reveals that cohesin stably bound to chromosomes regulates cohesin-mediated loop extrusion, and possibly LE mediated by other SMC complexes, by acting as semi-permissive as well as a non-permissive barrier when supporting SCC. These regulations may be key determinants in order to ensure sister chromatid resolution and correct segregation.

## Acknowledgements

This research was supported by the European Research Council under the Horizon 2020 Program (ERC grant agreement 771813) to RK, Agence Nationale pour la Recherche (ANR-22-CE12-0013-01) to FB and RK. FB also received support from the Fondation ARC pour la Recherche sur le Cancer (ARCPJA2022060005240) and the Comité de l’Occitanie de la Ligue Nationale contre le Cancer. NB was supported by the Ministère de l’Enseignement Supérieur and la Ligue Nationale contre le Cancer and Le comité départementale de la Ligue contre le Cancer de la Moselle. CC was supported by Pasteur-Roux-Cantarini fellowship.

We thank all our colleagues from the laboratories régulation spatiale des génomes and organization du noyau for helpful comments on the manuscript, K. Nasmyth’s laboratory and B. Albert for strains and K. Shirahige for the Smc3 antibody.

## Authors Contributions

N. B., F. B., and C. C. performed the experiments and analysis with contributions from A.T. All authors designed the research, interpreted the data and wrote the manuscript.

## Supplemental Information

### Supplementary Figure

**Supplementary Figure S1.**
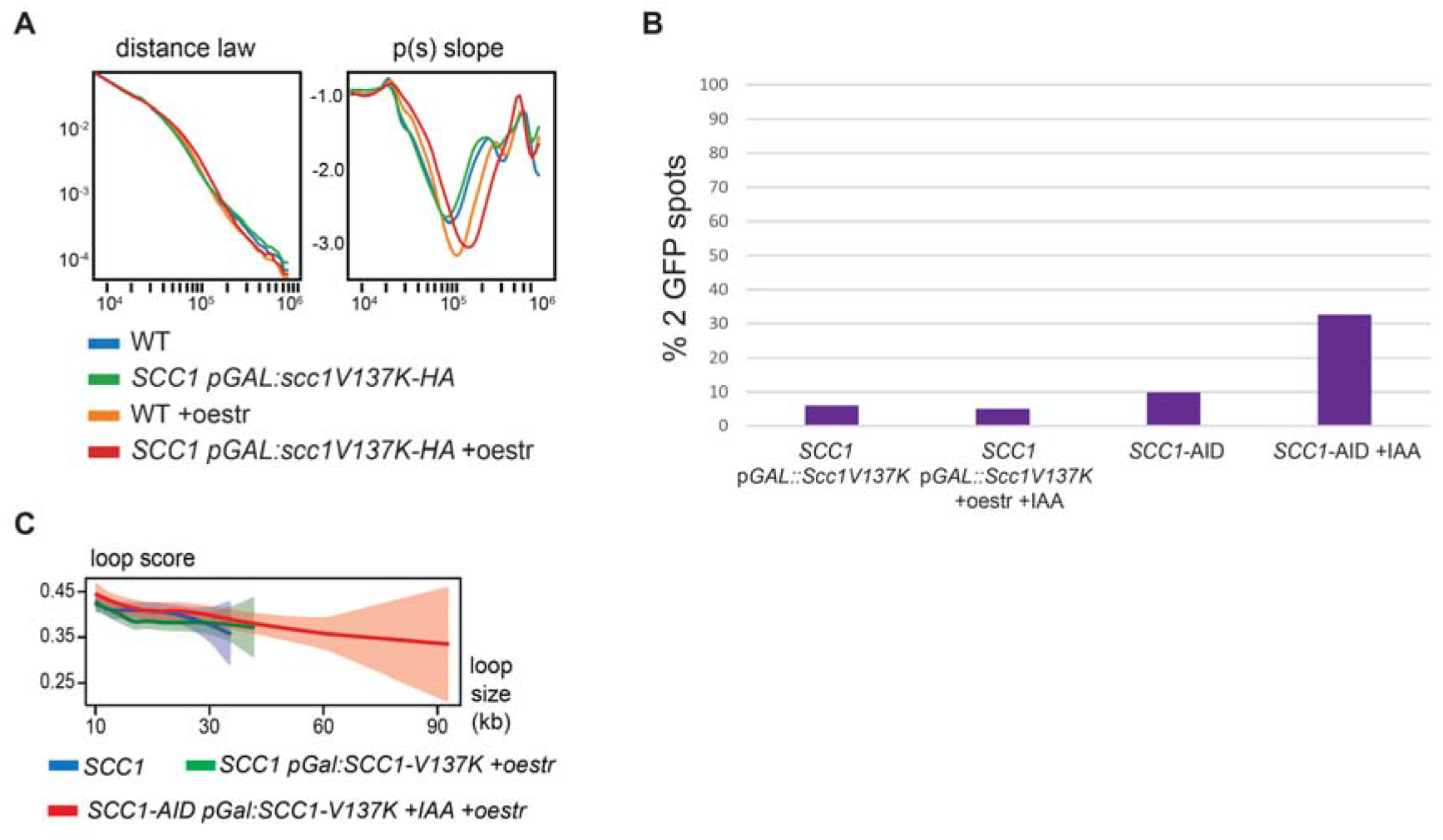
relative to Figure 2 and 3. **A)** Contact probability curves p(s) representing contact probability as a function of genomic distance (bp) and their respective derivative curves for WT cells and cells expressing *scc1-V137K* in presence of endogenous SCC1 before and after estradiol addition. **B)** Percentage of cells with paired or unpaired fluorescently labelled loci (leu2::TetR-GFP:LEU2, ChrXV 326K::tetO112::Kan) before and after *scc1-V137K* expression in presence of endogenous SCC1. Scc1-AID degradation by auxin was used as controlled of loss of cohesion. **C)** Average loop scores as a function of genomic distance quantified using Chromosight.

## Methods

### Data Availability

Raw data will be made available following peer-reviewed publication.

*Table S2* recapitulates the number of sequencing reads exploited for each condition tested.

*Reference genomes*

*S. cerevisiae* W303: gift from (O’Donnell et al., 2022);

*C. glabrata*: http://www.candidagenome.org/download/sequence/C_glabrata_CBS138/current/ Strains of this study are available from the corresponding authors.

### Code Availability

Programs involved in the study are listed below. They include custom made, open source programs from the Koszul lab:

HiCstuff (www.github.com/koszullab/hicstuff) version 3.0.1 Chromosight (www.github.com/koszullab/chromosight) v1.3.1 36 As well as programs published by others:

Bowtie2 (version 2.3.4.1 available online at http://bowtie-bio.sourceforge.net/bowtie2/) Samtools (version 1.9 available online at http://www.htslib.org/download/)

Bedtools (version 2.29.1 available online at https://bedtools.readthedocs.io/en/latest/content/installation.html)

Cooler (version 0.8.7-0.8.11 available online at https://cooler.readthedocs.io/en/latest/) bamCoverage (version 3.4.1 https://deeptools.readthedocs.io/en/develop/content/tools/bamCoverage.html)

### Yeast strains

Genotypes of the strains used in this study are described in Table S1.

Mutant *scc1-V137K* was constructed by directed mutagenesis using plasmid KN3752. The construction was verified by sequencing. Genomic integration was performed by homologous recombination in LEU2 locus.

### Media and growth conditions

Strains W303-1A, yNB33.1-8a, yNB77.1-2b and yNB78.1-15c were grown overnight at 30°C in 300mL of yeast peptone medium containing glucose (YPD): 1% bacto-peptone (Difco), 1% bacto yeast extract (Difco), and 2% glucose to attain 4,2.10^8 cells. Cells were synchronized in G1 by adding alpha-factor (Ref, Antibodies-online, ABIN399114) in the media every 30 minutes (1µg/mL final) during 2h30. After 1h of alpha-factor arrest auxin (2mM final) (Sigma-Aldrich, I3750) and estradiol (100nM final) (Sigma-Aldrich: E2758) were added to the media to respectively degrade Scc1-PK-AID and express SCC1 or Scc1V137K under control of pGAL1-10 promoter. Then cells were washed with fresh YP and released in YPD containing Nocodazole (Sigma-Aldrich, M1404-10MG) 2mM of auxin and 100nM of estradiol.

Strains yNB78.1-9a; yNB78.1-15c and BEN15, yNB84.1-17b and yNB81-5a were grown overnight in YPD at 30°C. Cells were synchronized by adding alpha-factor as describe above. Auxin was added to the media after 1h of alpha factor arrest to degrade Scc1-PK-AID and/or Cdc45-AID-FLAG. Then cells were washed and released in media supplemented with Nocodazole and auxin (2mM final). After 60min of Nocodazole arrest, estradiol (100nM final) was added to the media to express Scc1V137K during 90min.

### Flow cytometry

To verify cell cycle synchronization, arrest and absence of replication, the cells were analyzed using flow cytometry. For the different timepoints 1mL of cell culture were fixed in ethanol 70% and stored overnight at -20°C. Pellet was incubated with 50mM Tris-HCl (pH 7,5) and 5µL RNase A (10mg/mL) overnight at 37°C. Cells were pelleted and resuspended in 400µL of FACS buffer (1mg/mL propidium iodide (Fisher, P3566), Tris-HCl, NaCl, MgCl2) and incubated at 4°C during at least 30min. Then, cells were sonicated with 60% output for 10 seconds.

Flow cytometry was performed on a CytoFLEX S (Beckman Coulter). 1,000 events were counted with a sample flow rate of 30 µL/min and data were analyzed using CytExpert 2.4 software a CyFlow ML Analyzer (Partec).

### Proteins degradation analysis

A pellet of 10mL from experimental culture was frozen in liquid nitrogen and stored overnight at -20°C. Cell pellets were resuspended in 100 µL H2O and 20 µL trichloroacetic acid (Sigma-Aldrich, T8657) (TCA). Then cells were broken by glass beads at 4°C and precipitated proteins were resuspended in Laemmli buffer (100 mM DTT and Tris-HCl, pH 9,5). Proteins were extracted by cycles of 5 min heating at 80°C and 5 min vortexing at 4°C. After centrifugation, extracted proteins were collected and froze at -20°C.

Eluates were analyzed by SDS-page followed by western blotting with antibodies:

- Mouse anti Smc3-K113Ac (Beckouët et al., 2010) (Ab from K. Shirahige laboratory, clone H2) used at dilution 1:1000 for Western Blot
- Mouse anti-V5 tag (Bio-Rad, MCA1360) used at dilution 1:5000 for Western Blot
- Mouse anti-HA (Abcam, ab1424, 12CA5), used at dilution 1:1,000 for Western Blot
- Mouse anti-pgk1 (22C5D8) (Invitrogen, 459250) used at dilution 1:5000 for Western Blot
- Anti-mouse IgG, HRP conjugate (Promega, W4021) used at dilution 1:5000 for Western Blot

Blots were revealed using ChemiDoc Touch Imaging System: Image Lab 6.0.

### Calibrated Chip-Seq

Cells were grown exponentially to OD600 = 0.5. In triplicates, 15 OD600 units of S. cerevisiae cells were mixed with 3 OD600 units of C. glabrata to a total volume of 45 mL and fixed with 4mL of fixative solution (50 mM Tris-HCl, pH 8.0; 100 mM NaCl; 0.5 mM EGTA; 1 mM EDTA; 30% (v/v) formaldehyde) for 30 min at room temperature (RT) with rotation. The fixative was quenched with 2mL of 2.5M glycine (RT, 5 min with rotation). The cells were then harvested by centrifugation at 3,500 rpm for 3 min and washed with ice-cold PBS. The cells were then resuspended in 300 mL of ChIP lysis buffer (50 mM HEPES KOH, pH 8.0; 140 mM NaCl; 1 mM EDTA; 1% (v/v) Triton X-100; 0.1% (w/v) sodium deoxycholate; 1 mM PMSF; 2X Complete protease inhibitor cocktail (Roche)) and transfer in tubes 2mL containing glass beads before mechanical cells lysis. The soluble fraction was isolated by centrifugation at 2,000 rpm for 3min then transferred to sonication tubes and samples were sonicated to produce sheared chromatin with a size range of 200-1,000bp. After sonication the samples were centrifuged at 13,200 rpm at 4°C for 20min and the supernatant was transferred into 700µL of ChIP lysis buffer. 80µL (27µl of each sample) of the supernatant was removed (termed ‘whole cell extract [WCE] sample’) and store at -80°C. 5ug of antibody (anti-HA (Abcam ab1424, 12CA5)) was added to the remaining supernatant which is then incubated overnight at 4°C (wheel cold room). 50µL of protein G Dynabeads was then added and incubated at 4°C for 2h. Beads were washed 2 times with ChIP lysis buffer, 3 times with high salt ChIP lysis buffer (50mMHEPES-KOH, pH 8.0; 500 mM NaCl; 1 mM EDTA; 1% (v/v) Triton X-100; 0.1% (w/v) sodium deoxycholate;1 mM PMSF), 2 times with ChIP wash buffer (10 mM Tris-HCl, pH 8.0; 0.25MLiCl; 0.5% NP-40; 0.5% sodium deoxycholate; 1mM EDTA;1 mMPMSF) and 1 time with TE pH7.5. The immunoprecipitated chromatin was then eluted by incubation in 120µL TES buffer (50 mMTris-HCl, pH 8.0; 10 mM EDTA; 1% SDS) for 15min at 65°C and the supernatant is collected termed ‘IP sample’. The WCE samples were mixed with 40µL of TES3 buffer (50 mM Tris-HCl, pH 8.0; 10 mM EDTA; 3% SDS). ALL (IP and WCE) samples were de-cross-linked by incubation at 65°C overnight. RNA was degraded by incubation with 2µL RNase A (10 mg/mL) for 1h at 37°C. Proteins were removed by incubation with 10µL of proteinase K (18 mg/mL) for 2h at 65°C. DNA was purified by a phenol/Chloroform extraction. The triplicate IP samples were mixed in 1 tube and libraries for IP and WCE samples were prepared using Invitrogen TM ColibriCollibri TM PS DNA Library Prep Kit for Illumina and following manufacturer instructions. Paired-end sequencing on an Illumina NextSeq500 (2x35 bp) was performed. For analysis, Bowtie2 was used for two rounds of alignments, first on C. glabrata (CBS138) and then on S. cerevisiae allowing the generation of an alignment of IP and WCE that exclusively mapped on S. cerevisiae (an vice-versa for C. glabrata). The obtained SAM file was converted into a BAM file, sorted and indexed using Samtools. ChIP-seq profiles were then normalised by the number of million sequences and converted into BigWig using bamCoverage. Profiles were multiplied by the ORi factor (WCEglabrataIPcerevisiae / WCEcerevisiaeIPglabrata, in which WCE glabrata and IPglabrata correspond to the number of paired reads that mapped uniquely on C. glabrata genome and same for S. cerevisiae reads) using Integrated Genome Browser.

### Hi-C procedure and sequencing

Cell fixation with 3% formaldehyde (Sigma-Aldrich, Cat. F8775) was performed as described in Dauban et al. (2021). Quenching of formaldehyde with 300 mM glycine was performed at 4°C for 20 min. Hi-C experiments were performed using Arima Hi-C kit (Arima Genomics) with a double DpnII + HinfI restriction digestion. Briefly, samples were permeabilised by sequentially adding lysis buffer (15min at 4C), conditioning solution (10min at 62C) and stop solution 2 (15min at 37C) to the samples. DNA was digested using a mix of buffer A, DpnII, and Hinf1 (45min at 37C followed by 20min at 65C). DNA was repaired and biotin was added by adding a mix of buffer B and enzyme B for 45min at room temperature. DNA was re-ligated by adding a mix of buffer C and enzyme C during 15min at room temperature. Samples were then digested by protease and de-crosslinked by adding a mix of buffer D, enzyme D and buffer E during 30 min à 55°C followed by 90 min at 68°C. Samples were purified using AMPure XP beads (Beckman A63882), recovered in 120ul H2O and sonicated using Covaris (DNA 300bp). Preparation of the samples for paired-end sequencing on an Illumina NextSeq500 (2x35 bp) was performed using Invitrogen TM Collibri TM PS DNA Library Prep Kit for Illumina and following manufacturer instructions.

### Processing of the reads, computation of contact matrices, and generation of contact maps

Reads were aligned and the contact data processed using Hicstuff (https://github.com/koszullab/hicstuff). Briefly, pairs of reads were aligned iteratively and independently using Bowtie2 in its most sensitive mode against the S. cerevisiae W303 reference genome. Each uniquely mapped read was assigned to a restriction fragment. Quantification of pairwise contacts between restriction fragments was performed with default parameters: uncuts, loops and circularization events were filtered as previously described (Bastié et al, 2021).

PCR duplicates (defined as paired reads mapping at exactly the same position) were discarded. Contact maps from independent replicates were generated with the “*view*” function of Hicstuff, merged and normalized using the merge and balance functions of Cooler. Bins were set at 1, exp0.2 transformed, and rendered.

Final number of reads and corresponding experiments are listed in Table S2.

### Computation of the contact probability as a function of genomic distance

Computation of the contact probability as a function of genomic distance P(s) and its derivative have been determined using the “*distance law*” function of Hicstuff with default parameters, averaging the contact data of entire chromosome arms. P(s) from two independent replicates were determined and plotted.

### Loop detection and scoring with Chromosight

Chromosight 1.3.1 36 was used to call loops de novo from contact maps binned at 1 kb and balanced with Cooler. Matrices were subsampled to contain the same total number of contacts. De novo loop calling was computed using the “*detect*” mode of Chromosight, with minimum loop length set at 2kb, percentage undetected set at 25 and pearson correlation threshold set at 0.315. Loop strength was quantified for each loop using the quantify mode of Chromosight and the mean loop score was calculated for each condition. Loop pile-up of averaged 17kb windows were generated with Chromosight. De novo loop calling and scoring were performed on independent HiC replicates, where indicated, which were subsampled independently for each experiment, using the number of contacts of the least covered map or subsampling to 20M reads. Loop score is shown as mean ± standard deviation.

